# Enteropathogenic *E. coli* effector Map regulates the depletion of the tight junction proteins occludin and claudin-4 via cathepsin B and Rab13-mediated mechanisms

**DOI:** 10.1101/2024.02.12.579967

**Authors:** Anupam Mandal, Pangertoshi Walling, Kritika Kansal, Saima Aijaz

## Abstract

Infections by Enteropathogenic *E. coli* (EPEC) cause acute diarrheal disease in infants accounting for severe morbidity and mortality. One of the underlying causes of the disease is the break-down of the intestinal barrier maintained by the tight junctions (TJs). EPEC uses a type 3 secretion system to translocate more than twenty effectors into infected cells which disrupt several functions of the host cells. The effectors EspF, Map, EspG1/G2 and NleA have been reported to disrupt the TJs and cause the leakage of charged ions and uncharged molecules through the barrier. We have reported earlier that EspF and Map cause the depletion of TJ proteins claudin-1, claudin-4 and occludin through both transcriptional and post-transcriptional mechanisms. Here, we show that the EPEC effector Map modulates the lysosomal protease, cathepsin B to deplete claudins and occludin. Expression of mutant Map proteins that lacked the mitochondrial targeting sequence (MTS) completely restored the total levels of occludin and its localization at the TJs and partially restored claudin-4 levels and its junctional localization. We also identified a novel interaction of Map with the GTPase Rab13. As Rab13 has been reported to mediate the recycling of occludin to the plasma membrane, its interaction with Map has important implications for the loss of TJ integrity in EPEC pathogenesis. Occludin regulates the passage of water and uncharged solutes through TJs and Map may block its recycling to compromise the TJs thus causing excessive leakage through the barrier.

## INTRODUCTION

Enteropathogenic *E. coli* infection results in the rapid onset of diarrhea in infants in the developing world causing significant mortality (Hodges and Gill, 2010; Nataro and Kaper, 1998; Ochoa and Contreras, 2011). The infection is characterized by the disruption of the intestinal tight junction (TJ) barrier, defects in the sodium-glucose co-transporter 1 (SGLT-1), Na^+^/H^+^ exchanger isoform 3 (NHE3), Cl^-^/HCO_3_^-^ exchanger (also known as down-regulated in adenoma, DRA) and aquaporins (Croxen and Finlay, 2010; Guttman and Finlay, 2008; Viswanathan et al., 2009). The combined effect of these disruptions results in the excessive leakage of solutes and ions through the intestinal layer as well as reduced absorption of nutrients by the enterocytes (Vallance and Finlay, 2000; Viswanathan et al., 2009). EPEC uses a type 3 secretion system (T3SS) to translocate at least 20 effector proteins into the infected enterocytes (Dean and Kenny, 2009; Dean et al., 2005; Wong et al., 2011). Once inside the host cell, these effector proteins target multiple cell organelles and disrupt their functions. Of the more than twenty EPEC effectors identified so far, four effectors have been reported to disrupt the intestinal barrier. These include EspF, Map, EspG1/G2 and NleA (Dean and Kenny, 2004; Glotfelty et al., 2014; McNamara et al., 2001; Philpott et al, 1996; Singh and Aijaz, 2016; Singh et al., 2018; Thanabalasuriar et al., 2010; Ugalde-Silva et al., 2016). Extensive studies to unravel the molecular basis of EPEC pathogenesis have been hampered by the absence of a suitable animal model that mimics human infection. In the absence of a human model, earlier studies have mostly relied on a mouse model of infection as well as cell culture models (Law, et al., 2013; Savkovic, et al., 2005). These studies have highlighted the important roles of the EPEC effectors EspF and Map in mediating most of the disruptions in the host cell functions (Dean and Kenny, 2004; Dean et al., 2006; Holmes et al., 2010). For example, EspF inactivates the sodium hydrogen exchanger 3 (NHE3) and SGLT-1 while Map inactivates SGLT-1, Na^+^/H^+^ exchanger regulatory factor-1/-2 (NHERF-1/-2) (Dean et al., 2006; Hodges et al., 2008; Martinez et al., 2010; Simpson et al., 2006). In the intestine, the paracellular space between the epithelial cells is sealed by the TJs which selectively regulate the passage of ions and solutes through this space (Krug et al., 2014; Zihni et al., 2016). The TJ complex comprises transmembrane proteins such as occludin and claudins which regulate the passage of uncharged and charged molecules through the junctional space, respectively. The TJ proteins are frequently targeted by pathogens to disrupt the epithelial barrier (Croxen and Finlay, 2010). Both EspF and Map have been reported to cause extensive damage to the intestinal TJ barrier (Canil et al., 1993; Ma et al., 2006). Our earlier studies have shown that TJ disruption by EspF and Map is not only a consequence of the displacement of the membrane proteins from the TJs but also the significant depletion of their total levels in the cells (Singh, et al., 2018). This depletion was found to be caused by (i) the decrease in the transcripts of *occludin, claudin-1* and *claudin-4* and (ii) the decrease in the levels of the corresponding proteins by post transcriptional regulation (Singh, et al., 2018). We showed, for the first time, that the depletion of TJ proteins was partially reversed by inhibiting the lysosomes in cells expressing EspF or Map (Singh, et al., 2018). In continuation of that work, we wanted to further assess the involvement of the host cell lysosomes in the depletion of TJ proteins in response to the EPEC effector Map. Map is a 203 amino acid protein translocated into the infected cells via the T3SS (Dean and Kenny, 2009). Map harbors a mitochondrial targeting sequence (MTS) at the N-terminus between amino acid residues 1-44, a GTPase domain containing a conserved WxxxE motif between residues 74-78 and a PDZ Class1-binding domain containing the TRL motif at the C-terminus between amino acid residues 201-203 (Dean and Kenny, 2009). The N-terminal MTS of Map targets it to the mitochondria where it disrupts mitochondrial functions with the mitochondrial toxicity region (MTR) located between residues 101-152 playing an important role in altering the mitochondrial morphology (Papatheodorou, et al., 2006). The GTPase domain allows Map to act as a guanine nucleotide exchange factor for the activation of Cdc42 at the plasma membrane which regulates the formation of transient filopodia in the infected cells during the early stages of infection (Huang, et al., 2009). Activation of Cdc42 at the plasma membrane is mediated by the interaction between the TRL residues of Map and the PDZ domain of the ERM-binding phospho-protein 50 (Ebp50), also called NHERF-1, in an actin-dependent manner (Orchard et al., 2012; Simpson, et al., 2006). In order to study the role of the host cell lysosomes in the Map-mediated depletion of TJ proteins, we treated cells expressing GFP-tagged Map with lysosomal inhibitors and assessed the effect of this treatment on the total levels of the TJ proteins. We report here that the lysosomal cathepsins are involved in Map-mediated depletion of TJ membrane proteins. Further, by generating targeted deletions within Map, we have identified domains within Map that mediate the depletion of the TJ proteins as well as interactions with host proteins that regulate endocytosis such as caveolin-1 and Rab13.

## RESULTS AND DISCUSSION

We first reported the involvement of the lysosomes in the EspF- and Map-mediated depletion of TJ proteins wherein we found that treatment of cells with chloroquine caused a significant increase in the total levels of claudin-4 and occludin and to a lesser extent claudin-1 in cells expressing EspF and Map (Singh, et al., 2018). This increase was more evident in cells expressing Map (Singh, et al., 2018). Therefore, we further investigated the role of the lysosomal proteases in Map-mediated depletion of TJ proteins. Earlier studies have indicated that some TJ transmembrane proteins are targets of lysosomal cathepsins in different disease conditions. For example, the intestinal barrier breakdown in hemorrhagic shock is reported to be caused by the degradation of claudin-3 and occludin by the lysosomal cysteine protease, Cathepsin B and the inhibition of Cathepsin B prevented the degradation of these proteins (Klein, et al., 2017). Inhibition of another lysosomal cysteine protease, Cathepsin S has been reported to reverse TGF-β-induced epithelial-mesenchymal transition, restore the turnover of TJ proteins and prevent invasive growth in glioblastoma cells (Wei, et al., 2021). In retinal pigment epithelial cells, increased exposure to recombinant Cathepsin B disrupted the junctional localization of occludin and ZO-1 (Hadrian, et al., 2019). Further, in mouse models of induced and spontaneous colitis, treatment with mannose prevented the intestinal barrier damage by inhibiting Cathepsin B (Dong, et al., 2022).

To investigate if the depletion of claudin-1, claudin-4 and occludin in cells expressing Map was mediated by the lysosomal cathepsins, we treated the cells with E-64, an irreversible inhibitor of cathepsin B, H, and L (Barrett, et al., 1982) or CA-074 methyl ester (CA-074Me), a cell-permeable analog of CA-074 which irreversibly inhibits intracellular cathepsin B (Buttle, et al., 1992). Stable cell lines expressing either the AcGFP vector alone, AcGFP-Tir or AcGFP-Map were grown in individual wells of a 6 well plate until fully confluent and treated with E-64 (30μM) or CA-074Me (20μM) for 18 hours after which the total cell lysates were prepared and analyzed by western blotting (Figure 1). Treatment of cells with E-64 increased the total levels of claudin-1 to ∼0.87 and ∼0.89 fold and claudin-4 to ∼0.63 and ∼0.53 fold in the two clones expressing AcGFP-Map with respect to cells expressing AcGFP vector (Figure 1A, B). But the levels of occludin recovered significantly increasing to ∼0.94 and ∼1.38 fold with respect to cells expressing AcGFP vector after E-64 treatment (Figure 1A, B). Statistical differences were calculated by comparing fold changes between untreated and treated samples for each cell line as well as fold changes between each treated sample and AcGFP treated control samples. E-64 treated cell lines expressing Map (Map-1 and Map-2) showed no statistical difference from E-64 treated AcGFP cells in total occludin levels (Figure 1). No change was seen in the total levels of these proteins in cells expressing AcGFP-Tir which was used as a control cell line expressing Tir (<u>T</u>ranslocated intimin receptor), another EPEC effector which has no effect on TJ disruption (Singh, et al., 2018) We also used the cell-permeable, intracellular cathepsin B inhibitor, CA-074Me. Treatment of the above cell lines with CA-074Me increased the total levels of claudin-1 to ∼0.76 and ∼0.74 fold and claudin-4 to ∼0.64 and ∼0.72 fold. Again, the total levels of occludin in Map-expressing cells increased to ∼0.91 and ∼0.94 fold with respect to AcGFP cells treated with CA-074Me (Figure 1 B, C). E-64 and CA-074Me treatment resulted in a maximum increase in the levels of occludin, suggesting that Map modulates cathepsin B to cause the depletion of occludin in cells. The recovery seen in the total levels of claudin-1 and claudin-4 was also significant but not comparable to control cells indicating that other cathepsins may be involved in their depletion. A recent study has reported that EPEC caused the secretion of cathepsin D and β-hexosaminidase into the extracellular space and the aberrant appearance of the lysosomal membrane protein Lamp1 on the plasma membrane of infected cells and EspF and Map were attributed to play a role in this (Shtuhin-Rahav, et al., 2023). Our study provides further evidence that Map hijacks the lysosomes to disrupt the functions of the TJs by modulating cathepsin B. Next, we examined if this recovery in total levels of claudin-1, claudin-4 and occludin correlated with the recovery of these proteins at the plasma membrane. Cells expressing AcGFP vector alone, AcGFP-Tir or AcGFP-Map were grown on glass cover slips, treated with E-64 and CA-074Me and labeled with antibodies against claudin-1, claudin-4 and occludin. The cellular localization of these proteins in labeled cells was examined by immunofluorescence assays. As shown in Figure 2, the treatment with E-64 and CA-074Me did not increase the levels of these proteins at the TJs. However, their levels increased in the cytoplasm (Figure 2 A-C). Even for occludin, whose total levels were restored to the level of control cell lines in western blots (Figure 1), only discontinuous, patchy localization was seen at the TJs in cells treated with CA-074Me with most of the protein accumulating in the cytoplasm (Figure 2 C). This suggested that even when the turnover of these proteins was restored by cathepsin B inhibition, they were not transported to the TJs as was evident by their increased accumulation in the cytoplasm. Based on this data, we speculated that (i) Map delocalizes the existing TJ proteins and targets them to the lysosomes for degradation and (ii) Map also prevents the trafficking of the newly synthesized claudin-1, claudin-4 and occludin to the plasma membrane suggesting the involvement of the host protein transport/endocytosis machinery. In order to investigate this further, we generated targeted N-terminal and C-terminal deletions within Map to generate mutant proteins that (i) lacked the MTS located between amino acid residues 1-44 (MapΔ44), (ii) lacked the MTS and the GTPase domain containing the conserved WxxxE motif between residues 74-78 (MapΔ78) and (iii) lacked the TRL motif at the C-terminus between amino acid residues 201-203 (MapΔTRL). MDCK cell lines stably expressing these N-terminal GFP-tagged truncated proteins were generated and examined for their effect on the depletion of TJ proteins (Figure 3 A, B, C). Fold changes in the levels of TJ proteins in cell lines expressing different Map constructs were calculated with respect to cell lines expressing AcGFP vector alone. We observed that the increase in the total levels of claudin-1 in cells expressing truncated Map proteins was minimal. Cell lines expressing full length Map showed claudin-1 levels to be ∼0.45 fold and cell lines expressing MapΔTRL, MapΔ44 and MapΔ78 exhibited little recovery with claudin-1 expression levels of ∼0.39, ∼0.55 and ∼0.54 fold, respectively as compared to cell lines expressing AcGFP vector alone. For claudin-4, a significant increase of ∼0.72 fold was seen in cell lines expressing MapΔ44 (Figure 3 B, C) while cells expressing full length Map, MapΔTRL and MapΔ78 exhibited claudin-4 levels of ∼0.13, ∼0.2 and ∼0.4 fold respectively. Similarly, maximum recovery was observed in the levels of occludin in cell lines expressing MapΔ44 where the total levels of occludin were ∼1 fold which was similar to control cell lines (Figure 3 B, C). Occludin levels increased to ∼0.6 fold in cells expressing MapΔTRL. The levels of occludin in cells expressing full length Map and cells expressing MapΔ78 were ∼0.08 and ∼0.28 fold, respectively. This data suggests that the N-terminus MTS region of Map regulates the depletion of claudin-4 and occludin as its removal (in MapΔ44 construct) restored the levels of occludin completely and that of claudin-4 significantly. As cells expressing MapΔTRL also partially restored the levels of occludin, it appears that the C-terminus of Map has an as yet unidentified role in the recovery of occludin. Intriguingly, the MapΔTRL construct contains both the MTS and the mitochondrial toxicity region, yet the increase in the level of occludin in MapΔTRL cell lines was significant as compared to cells expressing full length Map. Immunofluorescence data showed a complete recovery of occludin at the TJs in cells expressing MapΔ44 and a partial junctional restoration of claudin-4 in MapΔ44 cells (Figure 4). However, cell expressing MapΔTRL, did not show any localization of occludin at the TJs (Figure 4 C) even though the western blotting data showed a significant increase in the total levels of occludin. The TRL domain of Map mediates its interaction with Ebp50, which in turn binds to the actin-binding protein ezrin, resulting in the activation of Cdc42 at the plasma membrane (Orchard et al., 2012). We speculated that the MTS domain of Map may cause the cathepsin B-mediated depletion of occludin and the TRL domain may be required for the trafficking of occludin to the plasma membrane.

**Figure 1:**
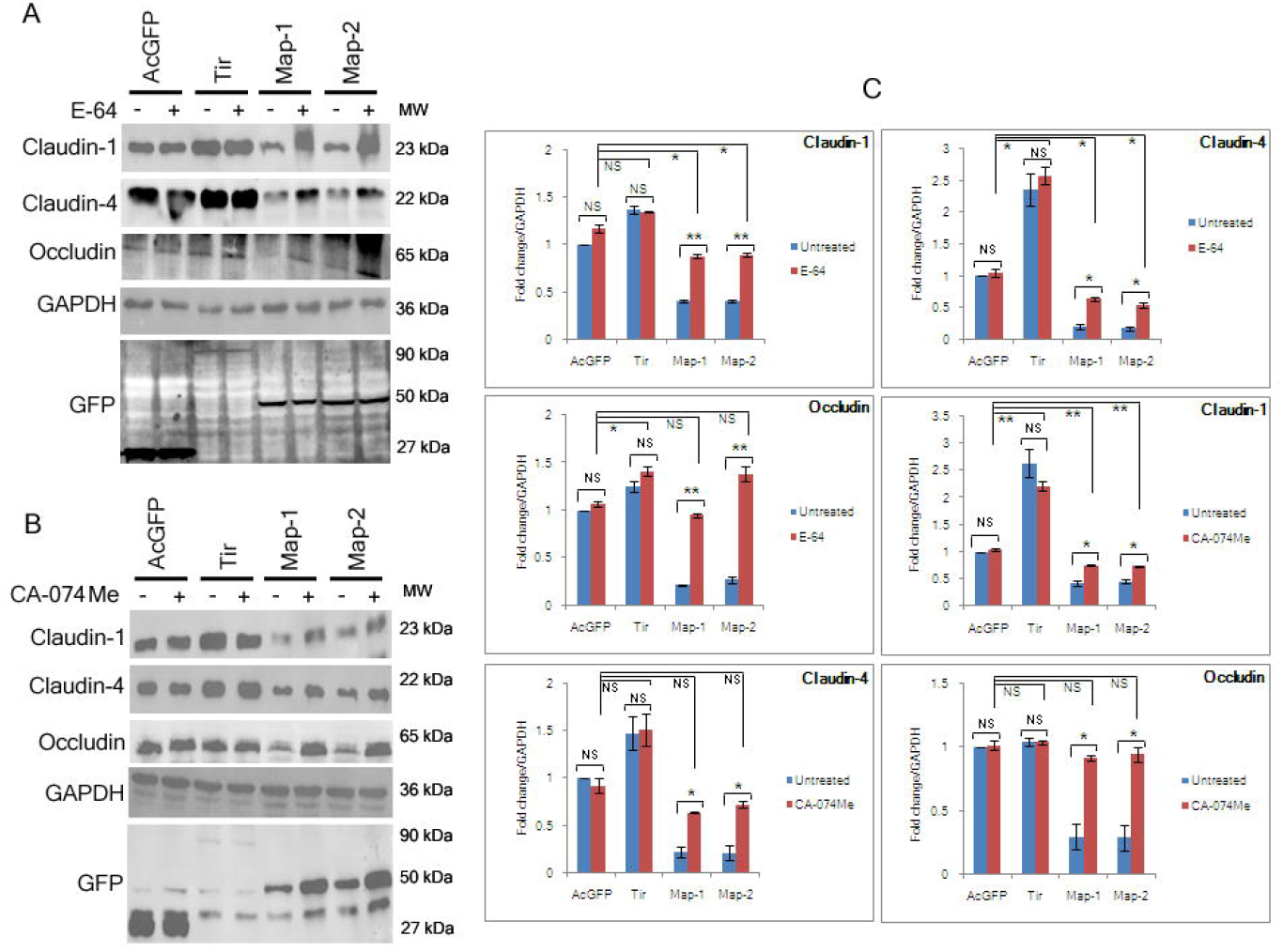
Effect of E-64 and CA-074Me treatment on the total levels of TJ proteins in cell lines expressing Map. Stable cell lines expressing AcGFP vector (labeled AcGFP), GFP-Tir (labeled Tir) and two independent cell lines expressing GFP-Map (labeled Map-1 and Map were treated with E-64 (A) and CA-074 (B) for 18 hours and cell lysates were prepared and subjected to western blotting with the indicated antibodies. GAPDH was used as a loading control. (C) The band intensities on the blots were analyzed by ImageJ software. The chart shows fold changes which were calculated by comparing untreated Tir and Map lysates with untreated AcGFP cell lysates (normalized to 1) and treated Map and Tir lysates with treated AcGFP lysates. Treatment with E-64 and CA-074Me increased the total levels of claudin-1, claudin-4 and occludin in cell lines expressing GFP-Map. Data are represented as means ± s.e.m. from at least 4-6 independent experiments. Horizontal lines in chart represent statistical differences between groups. *P<0.05; **P≤0.008; NS non significant.

**Figure 2:**
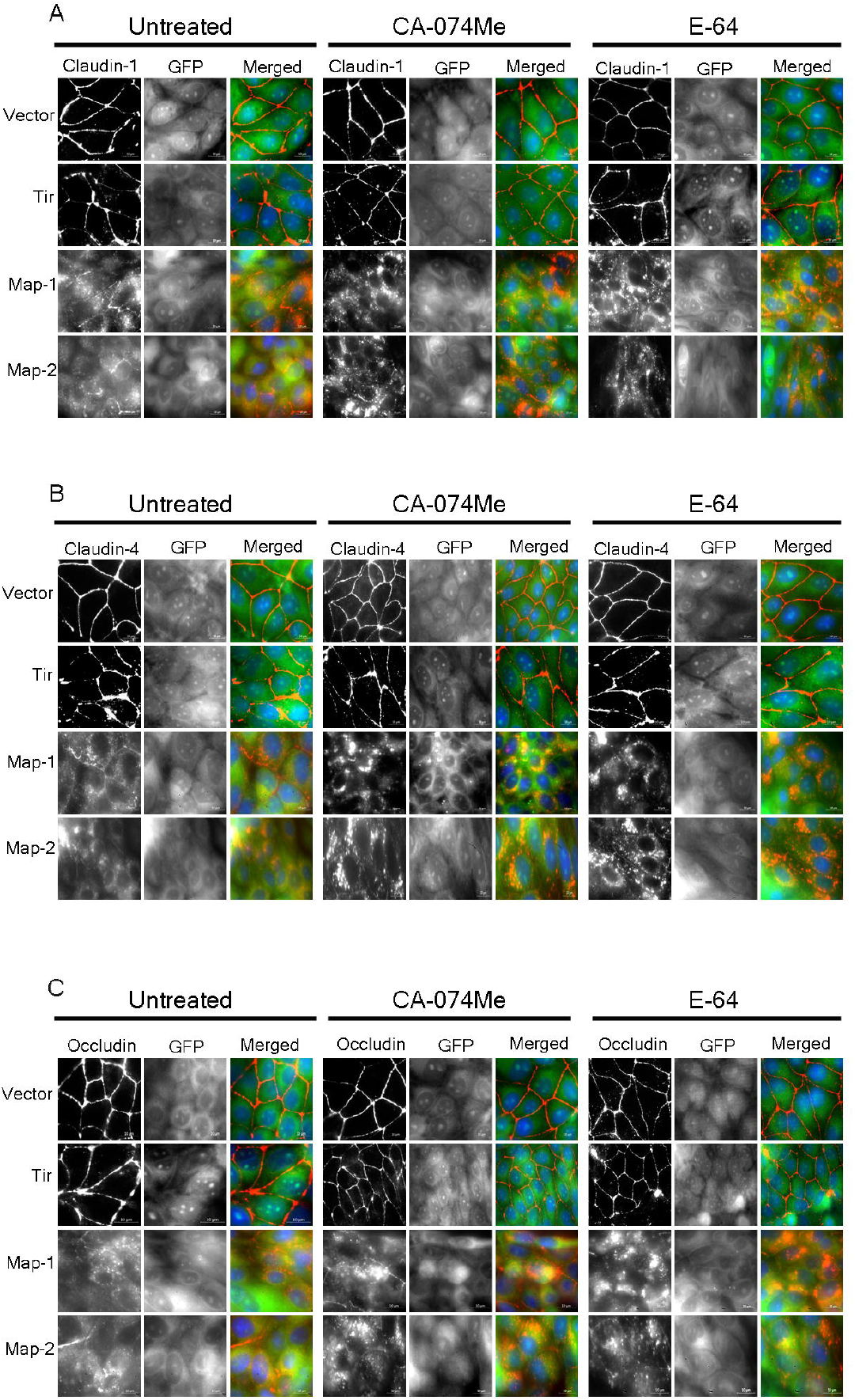
E-64 and CA-074Me treatment of cell lines expressing GFP-Map increases the accumulation of TJ proteins in the cytoplasm but not at the junctions. Stable cell lines expressing AcGFP vector, GFP-Tir and GFP-Map (Map1 and Map2 are two biological replicates) were cultured on glass cover slips until they were 90% confluent and then treated with E-64 (30μM) or CA-074Me (20μM) as described in the materials and methods section. Untreated cells were kept as controls. The cells were fixed with chilled methanol and then labeled with antibodies against the TJ proteins claudin-1 (A), claudin-4 (B) and occludin (C). The localization of TJ proteins in these cell lines was examined by fluorescence microscopy. Comparisons were made between cell lines expressing GFP-Map and control cell lines expressing either AcGFP vector or GFP-Tir. Increased accumulation of claudin-1, claudin-4 and occludin was observed in the cytoplasm of cells expressing GFP-Map but not in the control cell lines expressing AcGFP vector or GFP-Tir where these proteins were localized at the TJs. TJ proteins: red; GFP-tagged proteins: green; nucleus: blue. Scale bar: 10μm.

**Figure 3:**
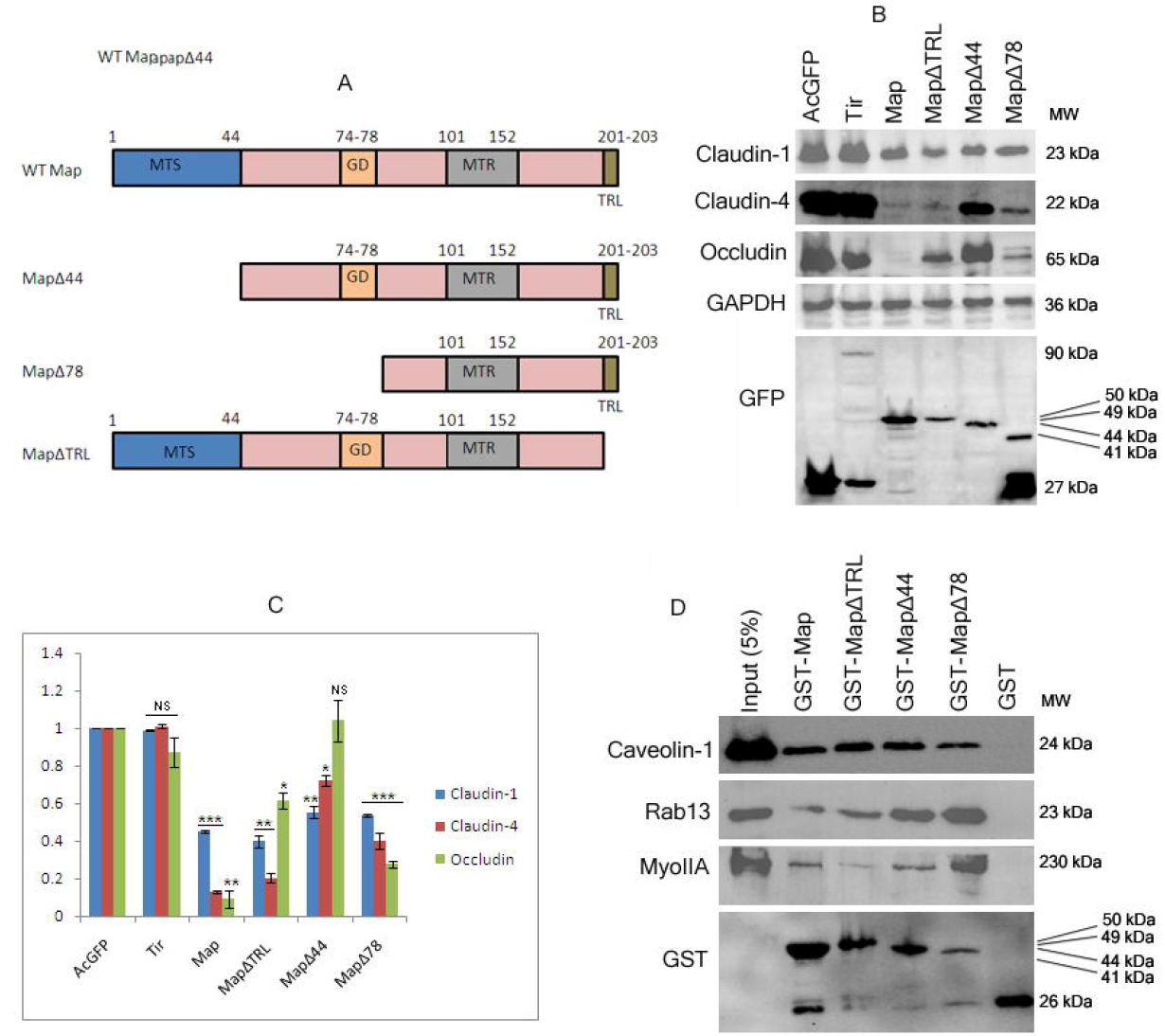
Role of Map domains in the depletion of TJ proteins and interactions with host endocytosis markers. (A) Schematic representation of Map showing the different domains. MTS: Mitochondrial targeting sequence; GD: GTPase domain containing WxxxE motif; MTR: Mitochondrial toxicity region; TRL: PDZ class 1-binding domain containing amino acid residues threonine, arginine and leucine, (B). Cell lysates derived from stable cell lines expressing AcGFP, GFP-Tir, GFP-Map, GFP-MapΔ44, GFP-MapΔ78 and GFP-MapΔTRL were analyzed by western blotting to identify domains within Map that mediate the depletion of TJ proteins. GAPDH was used as a loading control. (C) The band intensities were measured using ImageJ software. Fold changes in the expression of different TJ proteins in these cell lines were calculated with respect to lysates derived from cells expressing AcGFP vector (normalized to 1). Data are represented as means ± s.e.m. from at least 4 independent experiments. *P<0.05; **P<0.005; ***P<0.0005; NS non significant. Constitutive expression of truncated Map proteins increases the total levels of claudin-4 and occludin but not claudin-1 (D) Pull-down assays identified the interaction of different Map domains with caveolin-1, Rab13 and Myosin IIA.

**Figure 4:**
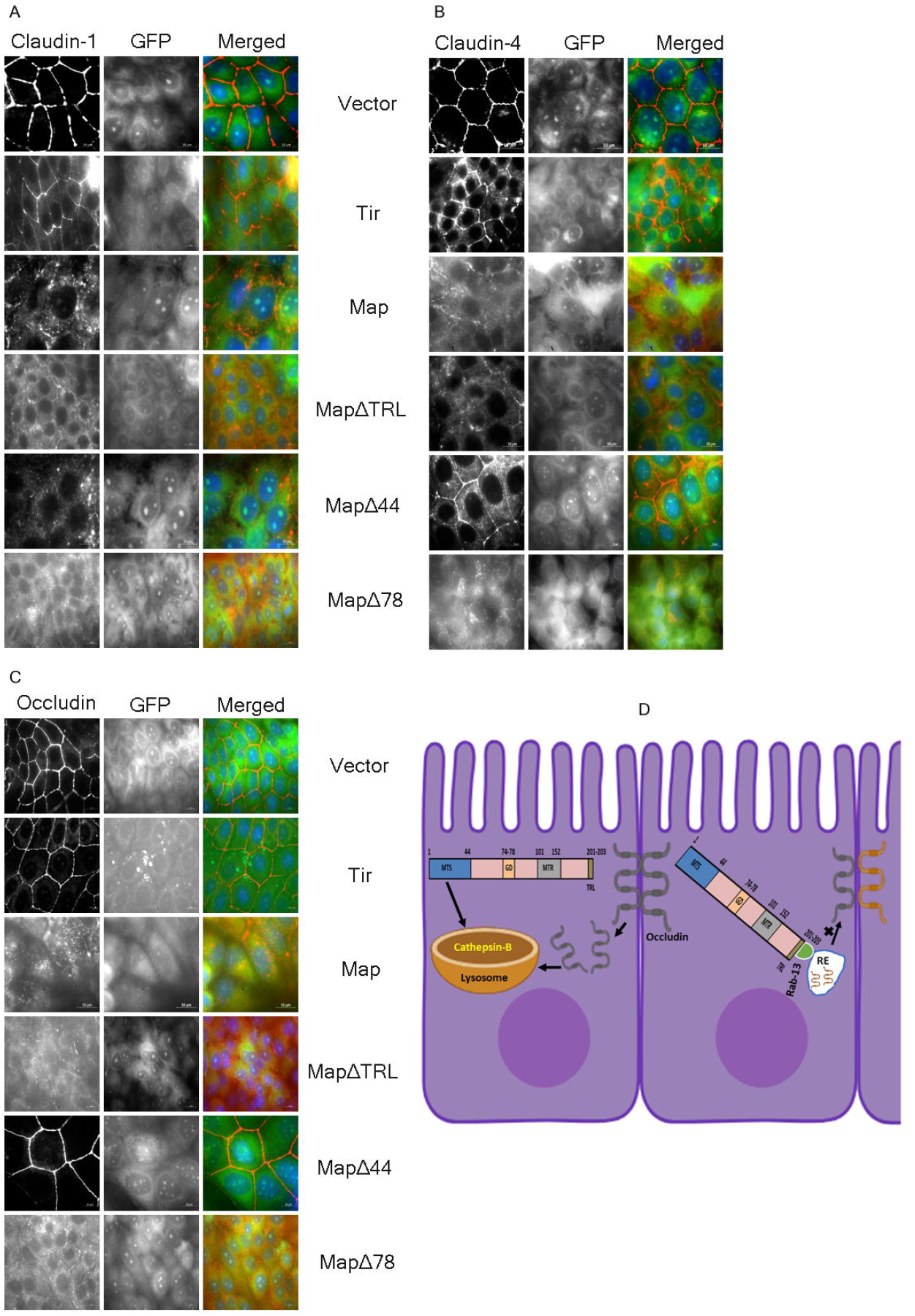
Localization of TJ proteins in cells expressing different Map domains. Stable cell lines expressing AcGFP vector alone, GFP-Tir, GFP-Map, GFP-MapΔ44, GFP-MapΔ78 or GFP-MapΔTRL were grown on cover slips and fixed with chilled methanol and labeled with antibodies against the TJ proteins, claudin-1 (A), claudin-4 (B) and occludin (C) to examine their cellular localization. Deletion of the first 44 residues of Map completely restored occludin at the TJs. Partial recovery of claudin-4 at the TJs was also observed although significant amount of claudin-4 was also localized in the cytoplasm of cells expressing GFP-MapΔ44. TJ proteins: red; GFP-tagged proteins: green; nucleus: blue. Scale bar: 10μm. (D) Schematic model showing the Map domains that likely link the lysosomes with the recycling endosome marker Rab13 to degrade existing junctional occludin/claudin-4 and prevent the trafficking of newly synthesized occludin/claudin-4 to the TJs. RE: recycling endosomes.

A recent study reported that Map causes the secretion of cathepsin D in the extracellular medium and the secretion of another lysosomal enzyme β-hexosaminidase was mediated via the GTPase domain and the MTR regions of Map (Shtuhin-Rahav, et al., 2023). Taken together, our data now suggests that Map may modulate the activity of multiple cathepsins and other lysosomal enzymes through different domains.

We further examined if the truncated Map proteins interacted with the host endocytosis machinery. Pull down assays were performed by immobilizing N-terminal GST-tagged Map proteins on glutathione sepharose beads and incubating them with MDCK cell lysates. All the truncated proteins were found to interact with caveolin-1 (Figure 3 D) to a similar extent. This suggests that Map may internalize plasma membrane proteins by caveolin-mediated endocytosis. In addition, we found that the GTPase Rab13 strongly interacted with MapΔ44 and MapΔ78 proteins suggesting that the residues 1-78 of Map were not involved in the interaction with Rab13. However, the efficiency of this interaction decreased when the TRL domain of Map was deleted (in MapΔTRL) which suggested that residues 79 to 203 likely mediate Rab13 interaction. Rab13 regulates the continuous recycling of occludin to and from the plasma membrane thus regulating TJs (Morimoto, et al., 2005) and a dominant active mutant of Rab13 was found to prevent this recycling (Morimoto, et al., 2005).Whether Map inhibits the GTPase activity of Rab13 to block the recycling of occludin to the TJs will form the basis of future investigations. EPEC has been reported to modulate Rab5a and Rab11 in the regulation of protein trafficking and polarity in an EspF-dependent manner (Kassa, et al., 2019; Tapia, et al., 2017). The interaction of Map with Rab13 reported here adds to a growing list of GTPases that are targeted by EPEC to disrupt host cell functions.

We earlier reported that EPEC Map interacts with myosin IIA (Singh, et al., 2018) and wanted to identify the domain of Map that interacts with Myosin IIA. Pull-down assays showed that MapΔTRL protein interacted only weakly with Myosin IIA (Figure 3, D). Myosin IIA is an important regulator of actin contractility which opens the TJ barrier (Zihni et al., 2016).Therefore, it is interesting to speculate that the TRL domain may serve to link the Ebp50-ezrin-actin complex with myosin IIA to cause actin contractility and contribute to the leakage of solutes across the TJ barrier in EPEC infection.

In conclusion, our data shows, for the first time, that Map depletes the TJ proteins via a cathepsin B-mediated pathway. Inhibition of cathepsin B increased the total levels of TJ proteins in the cytoplasm but not at the TJs. Truncated Map proteins that lacked the first 44 residues (MTS) were able to completely restore occludin and partially restore claudin-4 at the TJs pointing to a role of the MTS in their depletion. Additionally, we show that the GTPase Rab13 interacts with Map possibly through the TRL domain which may play an important role in the recycling of occludin to the TJs (Figure 4 D). The displacement of claudins and occludin is a critical event that causes increased permeability of ions and water/uncharged molecules, respectively through the TJs in EPEC diarrhea. Future studies on dissecting these interactions will aid in the development of therapies that can seal the TJ barrier in EPEC infections thus preventing the loss of life.

## MATERIALS AND METHODS

### Generation of plasmid constructs and recombinant Map proteins

The following primers containing restriction sites for EcoRI (at the 5’-end) and SalI (at the 3’-end) were used to generate full length and truncated *Map* constructs

Map forward: 5’-AAAAAGAATTCCCTTAAGATGGTTAGTCCAACGGCAATGGTA-3’

Map reverse: 5’-AAAAATCTAGAGTCGACCAGCCGAGTATCCTGCACATTGT-3’

MapΔ44-forward: 5’-AAAAAGAATTCCCTTAAGATGTCGAACCTTATGATTAATC-3’

MapΔ78-forward: 5’-AAAAAGAATTCCCTTAAGATGCAGATTACTTTTCTATCCA-3’

MapΔTRL-reverse: 5’-AAAAATCTAGAGTCGACATCCTGCACATTGTCTGCA-3’

PCR was performed by following standard protocols. Genomic DNA from Enteropathogenic *E. coli* O127:H6 strain E2348/69 was used as a template to generate *Map* constructs expressing full-length Map, MapΔTRL (lacking the C-terminal TRL residues), MapΔ44 (lacking residues 1 to 44) and MapΔ78 (lacking residues 1 to 78) proteins. The PCR products were cloned in pAcGFP1-C1 vector (for N-terminal GFP tag) or pGEX-4T-3 vector (for N-terminal GST tag) between the EcoRI and SalI sites of both vectors.

Additionally, GFP-tagged Tir (<u>T</u>ranslocated <u>i</u>ntimin <u>r</u>eceptor) construct was generated by using primers, containing restriction sites for BamHI (at 5’-end) and SalI (at 3’-end), shown below:

Tir forward: 5’-AAAAAGGATCCCTTAAGATGGCTATTGGTAACCTTGGT-3’

Tir reverse: 5’-AAAAATCTAGAGTCGACAACGAAACGTACTGGTCCCGG-3’

The resulting PCR products were cloned in pAcGFP1-C1 vector between the BglII and SalI sites. Tir was used as a control EPEC effector as it has no effect on the disruption of TJs.

Recombinant GST-tagged Map proteins were obtained by transforming BL21(DE3)pLysS cells with the plasmid constructs cloned in pGEX-4T-3 vector. Cultures were induced with 1mM IPTG and incubated overnight at 25ºC to obtain GST, full-length GST-Map, GST-MapΔ44, GST-MapΔ78 and GST-MapΔTRL proteins.

### Generation of stable cell lines

MDCK cells, maintained in Dulbecco’s Modified Eagle Medium (DMEM) supplemented with 10% Fetal Bovine Serum (FBS), were transfected with AcGFP1-C1 vector or the Map and Tir constructs cloned in pAcGFP1-C1 vector using Lipofectamine 3000 reagent (Thermo Fisher Scientific). Cells expressing AcGFP alone, full length Map, full length Tir and truncated Map proteins were selected by adding G418 (500 μg/ml) to the medium and incubated in a CO_2_ incubator at 37°C for 2-3 weeks. Subsequently, clones expressing AcGFP, GFP-Map, GFP-MapΔ44, GFP-MapΔ78, GFP-MapΔTRL and full length GFP-Tir were picked and propagated further. At least 10 clones were picked and analyzed for each cell line. Expression of all proteins was confirmed by western blotting with anti-GFP antibodies as well as by fluorescence microscopy. For each experiment, at least two independent clones (biological replicates) of Map were used.

### Immunofluorescence assays and microscopy

The cells expressing AcGFP vector alone, GFP-Tir, GFP-Map, GFP-MapΔ44, GFP-MapΔ78 and GFP-MapΔTRL were grown on glass cover slips until 90% confluent and then fixed with chilled methanol for 5 minutes followed by incubation in PBS for 5 minutes at room temperature. The cells were then incubated in blocking solution (containing 0.5% w/v BSA in PBS) for 30 minutes at room temperature followed by incubation with primary antibodies diluted 1:300 in solution containing 1XPBS, 0.5% BSA and 0.1% sodium azide for 2-4 hours at room temperature. The cover slips were then rinsed thrice with PBS, 0.5% BSA and 0.1% sodium azide. The cells on cover slips were then incubated with Cy3-conjugated anti-mouse secondary antibodies (Millipore) for 1 hour at room temperature. The cells were rinsed thrice and mounted in PBS, 0.5% BSA and 30% glycerol on glass slides. Images were obtained at 100X magnification on an ApoTome (Axiovert40 CFL, Zeiss). Primary antibodies used for labeling were claudin-1 (#374900, mouse monoclonal), claudin-4 (#329400, mouse monoclonal), occludin (#331500, mouse monoclonal), all purchased from Thermo Fisher Scientific. For labeling the nucleus, DAPI (4’,6-diamidino-2-phenylindole (Millipore) was used at 1:5,000 dilution.

### Preparation of total cell lysates

The total protein lysates were prepared from confluent cell lines expressing AcGFP vector alone, GFP-Map, GFP-MapΔ44, GFP-MapΔ78, GFP-MapΔTRL and GFP-Tir, grown on 6-well plates. The total cell lysates were prepared by adding 1X Laemmli buffer to the wells of culture plates followed by extraction through a 23-gauge needle several times. For CA-074Me and E-64 treatment, equal number of cells from cell lines expressing AcGFP vector, full-length GFP-Map, and GFP-Tir were seeded on a 6 well plate. At 90% confluency, the cells were treated with either E-64 (30 μM) or CA-074Me (20 μM) for 18 hours at 37°C and then rinsed with PBS. Total cell lysates, after treatment with E-64 or CA-074Me, were prepared as described above. The lysates were analyzed by electrophoresis on 12% SDS polyacrylamide gels followed by western blotting with primary antibodies at 1:1,000 dilution in blocking solution. The primary antibodies used were claudin-1, claudin-4, occludin and GFP (#A01388, rabbit polyclonal, GenScript). GAPDH (#AB0060, rabbit polyclonal, Biobharti Life Science; 1:10,000 dilution) was used as a loading control. Secondary antibodies used were HRP-conjugated anti-mouse or anti-rabbit antibodies (Millipore). After western blotting, the band intensities were analyzed by ImageJ software (https://imagej.net/ij) to obtain fold changes with respect to lysates derived from cell lines expressing AcGFP vector (normalized to 1).

### Pull-Down Assays

Lysates derived from BL21(DE3)pLysS cells expressing GST, full-length GST-Map, GST-MapΔ44, GST-MapΔ78 and GST-MapΔTRL proteins were clarified by centrifugation and incubated with glutathione sepharose beads overnight at 4ºC with constant rotation. The glutathione sepharose beads bound to GST, GST-Map, GST-MapΔ44, GST-MapΔ78 and GST-MapΔTRL were centrifuged, washed with lysis buffer containing 1X PBS, 1mM DTT, 1 mM PMSF and 0.5% Triton X-100 three times and then incubated with lysates derived from wild type MDCK cells for 12-14 hours at 4ºC on a rotary shaker. MDCK cell lysates were prepared by growing the cells on 100mm culture plates until fully confluent after which 1mL of pull-down buffer (1X PBS, 1mM DTT, 1mM PMSF and 0.5% Triton X-100) was added. Subsequently, the cells were harvested by scraping from the culture plates and passed through a 21-gauge needle several times and then incubated on ice for 30 minutes. The lysates were centrifuged and the supernatant was incubated with glutathione sepharose beads bound to the GST proteins mentioned above overnight at 4°C with rotation. The beads were centrifuged, washed three times with cell lysis buffer, mixed with equal volumes of 2X SDS gel loading dye, resolved on 12% SDS polyacrylamide gels and analyzed by western blotting. The primary antibodies used were caveolin-1 (#3267S, Rabbit monoclonal, Thermo Fisher Scientific), Rab13 (#PA5-52039, Rabbit monoclonal, Thermo Fisher Scientific), MyosinIIA ((#3403, rabbit polyclonal, Cell Signaling Technology) and GST (#GE27-4577-01, goat polyclonal, Sigma Aldrich; used at 1:10,000 dilution). HRP-conjugated anti-goat and anti-rabbit secondary antibodies (Millipore) were used at 1:10,000 dilution.

### Statistical Analysis

All experiments were performed three to four times with two independent clones (biological replicates) expressing full length GFP-Map or truncated Map proteins. Comparisons were made between cell lines expressing Map proteins and cell lines expressing AcGFP vector using unpaired t-tests. Differences between two groups were considered significant at p-values < 0.05.

## Acknowledgements

A.M., P.W. and K.K. acknowledge the receipt of research fellowships from University Grants Commission (A.M.), Council of Scientific and Industrial Research (P.W.) and Department of Biotechnology, India, (K.K.).

## Author contributions

Conceptualization: S.A.; Validation: A.M., S.A.; Formal analysis: S.A.; Investigation: A.M., P.W., K.K.; Resources: S.A.; Data curation: A.M., S.A.; Writing - original draft; S.A.; Writing - review & editing: S.A.; Visualization: S.A.; Supervision: S.A.; Project administration: S.A.; Funding acquisition: S.A.

## Data availability

All relevant data can be found within the article and its supplementary information.

## References

Barrett, A. J., Kembhavi, A. A., Brown, M. A., Kirschke, H., Knight, C.G., Tamai, M. and Hanada, K. (1982). “L-trans-Epoxysuccinyl-leucylamido(4-guanidino)butane (E-64) and its analogues as inhibitors of cysteine proteinases including cathepsins B, H and L.” Biochem. J. 201(1), 189–98.

Buttle, D. J., Murata, M., Knight, C. G. and Barrett, A. J.(1992). CA074 methyl ester: a pro inhibitor for intracellular cathepsin B. Arch. Biochem. Biophys. 299(2), 377–80.

Canil, C., Rosenshine, I., Ruschkowski, S., et al. (1993). Enteropathogenic Escherichia coli decreases the transepithelial electrical resistance of polarized epithelial monolayers. Infect. Immun. 61(7), 2755–2762.

Croxen, M. A. and Finlay, B. B. (2010). Molecular mechanisms of Escherichia coli pathogenicity. Nat. Rev. Microbiol. 8(1), 26–38.

Dean, P. and Kenny, B. (2009). The effector repertoire of enteropathogenic E. coli: ganging up on the host cell. Curr. Opin. Microbiol. 12, 101–109.

Dean, P., Maresca, M. and Kenny, B. (2005). EPEC’s weapons of mass subversion. Curr. Opin. Microbiol. 8(1), 28–34.

Dean, P., Maresca, M., Schüller, S., et al. (2006). Potent diarrheagenic mechanism mediated by the cooperative action of three enteropathogenic Escherichia coli-injected effector proteins. Proc. Natl. Acad. Sci. USA 103(6), 1876–1881.

Dean, P. and Kenny, B. (2004). Intestinal barrier dysfunction by enteropathogenic Escherichia coli is mediated by two effector molecules and a bacterial surface protein. Mol. Microbiol. 54, 665–675.

Dong, L., Xie, J., Wang, Y., Jiang, H., Chen, K., Li, D., Wang, J., Liu, Y., He, J., Zhou, J., Zhang, L., Lu, X., Zou, X., Wang, X. Y., Wang, Q., Chen, Z. and Zuo, D. (2022). Mannose ameliorates experimental colitis by protecting intestinal barrier integrity. Nat. Commun. 13(1), 4804.

Glotfelty, L. G., Zahs, A., Hodges, K., et al. (2014). Enteropathogenic E. coli effectors EspG1/G2 disrupt microtubules, contribute to tight junction perturbation and inhibit restoration. Cell. Microbiol. 16, 1767–1783.

Guttman, J. A. and Finlay, B. B. (2008). Subcellular alterations that lead to diarrhea during bacterial pathogenesis. Trends Microbiol. 16(11), 535–54.

Hadrian, K., Lever, M., Schlatt, S., Wistuba, J., Thanos, S., Rahmann, S., Klein-Hitpass, L. and Böhm, M. (2019). Expression and function of Cathepsin B in the retinal pigment epithelium. Invest. Ophthalmol. Vis. Sci. 60(9), 2383.

Hodges, K. and Gill, R. (2010). Infectious diarrhea. Gut Microbes 1:1, 4–21.

Hodges, K., Alto, N. M., Ramaswamy, K., et al.(2008). The Enteropathogenic Escherichia coli effector protein EspF decreases sodium hydrogen exchanger 3 activity. Cell. Microbiol. 10(8), 1735–1745.

Holmes, A., Muhlen, S., Roe, A. J., et al. (2010). The EspF effector, a bacterial pathogen’s swiss army knife. Infect. Immun. 78, 4445–4453.

Huang, Z., Sutton, S. E., Wallenfang, A. J., et al. (2009). Structural insights into host GTPase isoform selection by a family of bacterial GEF mimics. Nat. Struct. Mol. Biol. 16(8), 853–860.

Kassa, E. G., Zlotkin-Rivkin, E., Friedman, G., et al. (2019). Enteropathogenic Escherichia coli remodels host endosomes to promote endocytic turnover and breakdown of surface polarity. PLoS Pathog. 15(6), e1007851.

Klein, N.P., Courelli, A. and Schmid-Schoenbein, G. (2017). Cathepsin B Mediated Degradation of Jejunal Epithelial Tight Junction Proteins during the Early Stage of Hemorrhagic Shock. The FASEB Journal 31, 680.5.

Krug, S. M., Schulzke, J. D. and Fromm, M. (2014). Tight junction, selective permeability, and related diseases. Semin. Cell. Dev. Biol. 36, 166–176.

Law, R. J., Gur-Arie, L., Rosenshine, I., et al. (2013). In vitro and in vivo model systems for studying enteropathogenic Escherichia coli infections. Cold Spring Harb. Persp. Med. 3(3), a009977.

Ma, C., Wickham, M. E., Guttman, J. A., et al. (2006). Citrobacter rodentium infection causes both mitochondrial dysfunction and intestinal epithelial barrier disruption in vivo: role of mitochondrial associated protein (Map). Cell. Microbiol. 8, 1669–1686.

Martinez, E., Schroeder, G. N., Berger, C.N., et al.(2010). Binding to Na+/H+ exchanger regulatory factor 2 (NHERF2) affects trafficking and function of the enteropathogenic Escherichia coli type III secretion system effectors Map, EspI and NleH. Cell. Microbiol. 12(12), 1718–1731.

McNamara, B. P., Koutsouris, A., O’Connell, C. B., et al. (2001). Translocated EspF protein from enteropathogenic Escherichia coli disrupts host intestinal barrier function. J. Clinic. Investig. 107, 621–629.

Morimoto, S., Nishimura, N., Terai, T., Manabe, S., Yamamoto, Y., Shinahara, W., Miyake, H., Tashiro, S., Shimada, M. and Sasaki, T. (2005). Rab13 mediates the continuous endocytic recycling of occludin to the cell surface. J. Biol. Chem. 280, 2220–2228.

Nataro, J. P. and Kaper, J.B. (1998). Diarrheagenic Escherichia coli. Clin. Microbiol. Rev. 11, 142–201.

Ochoa, T. J. and Contreras, C. A. (2011) Enteropathogenic Escherichia coli infection in children. Curr. Opin. Infect. Dis. 24, 478–483.

Orchard, R. C. et al. (2012). Identification of F-actin as the dynamic hub in a microbial-induced GTPase polarity circuit. Cell 148, 803–815.

Papatheodorou, P., Domanska, G., Oxle, M., Mathieu, J., Selchow, O., Kenny, B. and Rassow, J. (2006). The enteropathogenic Escherichia coli (EPEC) Map effector is imported into the mitochondrial matrix by the TOM/Hsp70 system and alters organelle morphology. Cell. Microbiol. 8, 677–689.

Philpott, D. J., McKay, D. M., Sherman, P. M., et al. (1996). Infection of T84 cells with enteropathogenic Escherichia coli alters barrier and transport functions. Am. J. Physiol. 270, G634–645.

Savkovic, S. D., Villanueva, J., Turner, J. R., et al. (2005). Mouse model of enteropathogenic Escherichia coli infection. Infect. Immun. 73, 1161–1170.

Shtuhin-Rahav, R., Olender, A., Zlotkin-Rivkin, E., Bouman, E. A., Danieli, T., Nir-Keren, Y., Weiss, A. M., Nandi, I. and Aroeti, B. (2023). Enteropathogenic E. coli infection co-elicits lysosomal exocytosis and lytic host cell death. mBio. 14(6), e0197923.

Simpson, N., Shaw, R., Crepin, V. F., et al. (2006). The Enteropathogenic Escherichia coli type III secretion system effector Map binds EBP50/NHERF1: implication for cell signalling and diarrhoea. Mol. Microbiol. 60(2), 349–363.

Singh, A. P. and Aijaz, S. (2016). Enteropathogenic E. coli breaking the intestinal tight junction barrier. F1000Res. 4, 231.

Singh, A.P., Sharma, S., Pagarware, K., Siraji, R. A., Ansari, I., Mandal, A., Walling, P. and Aijaz, S. (2018). Enteropathogenic E. coli effectors EspF and Map independently disrupt tight junctions through distinct mechanisms involving transcriptional and posttranscriptional regulation. Sci. Rep. 8, 3719.

Tapia, R., Kralicek, S. E. and Hecht, G. A. (2017). EPEC effector EspF promotes Crumbs3 endocytosis and disrupts epithelial cell polarity. Cell. Microbiol. 19, e12757.

Thanabalasuriar, A., Koutsouris, A., Weflen, A., et al.(2010). The bacterial virulence factor NleA is required for the disruption of intestinal tight junctions by enteropathogenic Escherichia coli. Cell. Microbiol. 12(1), 31–41.

Ugalde-Silva, P., et al. (2016). Tight junction disruption induced by type 3 secretion system effectors injected by enteropathogenic and enterohemorrhagic Escherichia coli. Front. Cell. Infect. Microbiol. 6, 1162.

Vallance, B. A. and Finlay, B. B. (2000). Exploitation of host cells by enteropathogenic Escherichia coli. Proc. Natl. Acad. Sci. U S A. 97(16), 8799–8806.

Viswanathan, V. K., Hodges, K. and Hecht, G. (2009). Enteric infection meets intestinal function: how bacterial pathogens cause diarrhoea. Nat. Rev. Microbiol. 7(2), 110–119.

Wei, L., Shao, N., Peng, Y. and Zhou, P. (2021). Inhibition of Cathepsin S Restores TGF-β-induced Epithelial-to-mesenchymal Transition and Tight Junction Turnover in Glioblastoma Cells. J. Cancer 12(6), 1592–1603.

Wong, A. R., Pearson, J. S., Bright, M. D., et al.(2011). Enteropathogenic and enterohaemorrhagic Escherichia coli: even more subversive elements. Mol. Microbiol. 80(6), 1420–1438.

Zihni, C., Mills, C., Matter, K., et al. (2016). Tight junctions: from simple barriers to multifunctional molecular gates. Nat. Rev. Mol. Cell Biol. 17, 564–580.

